# Development of a transboundary model of livestock disease in Europe

**DOI:** 10.1101/2021.04.27.441716

**Authors:** Richard Bradhurst, Graeme Garner, Márk Hóvári, Maria de la Puente, Koen Mintiens, Shankar Yadav, Tiziano Federici, Ian Kopacka, Simon Stockreiter, Ivanka Kuzmanova, Samuil Paunov, Vladimir Cacinovic, Martina Rubin, Jusztina Szilágyi, Zsófia Szepesiné Kókány, Annalisa Santi, Marco Sordilli, Laura Sighinas, Mihaela Spiridon, Marko Potocnik, Keith Sumption

## Abstract

Epidemiological models of notifiable livestock disease are typically framed at a national level and targeted for specific diseases. There are inherent difficulties in extending models beyond national borders as details of the livestock population, production systems and marketing systems of neighbouring countries are not always readily available. It can also be a challenge to capture heterogeneities in production systems, control policies, and response resourcing across multiple countries, in a single transboundary model.

In this paper we describe EuFMDiS, a continental-scale modelling framework for transboundary animal disease, specifically designed to support emergency animal disease planning in Europe. EuFMDiS simulates the spread of livestock disease within and between countries and allows control policies to be enacted and resourced on per-country basis. It provides a sophisticated decision support tool that can be used to look at the risk of disease introduction, establishment and spread; control approaches in terms of effectiveness and costs; resource management; and post-outbreak management issues.

## 1. INTRODUCTION

An outbreak of notifiable livestock disease such as foot-and-mouth disease (FMD), rinderpest, highly pathogenic avian influenza (HPAI), peste des petits ruminants (PPR), African swine fever (ASF) or classical swine fever (CSF), can have serious socio-economic consequences for the afflicted country. In a multinational setting such as the European Union (EU) where there are high levels of trade and travel between member states, there is increased risk that a highly contagious livestock disease may silently cross borders via movements of presymptomatic infected livestock, contaminated livestock products or fomites (Beltran-Alcrudo et al., 2019). High impact and contagious livestock diseases that readily spread between countries are commonly referred to as Transboundary Animal Diseases (TADs) (Otte et al., 2004).

Whilst the detection and control of livestock diseases are largely national concerns, there is a strong case for international cooperation with respect to early detection systems, contingency planning, movement tracing systems, sharing of outbreak data, and coordination of control programs (Domenech et al., 2006; Martin et al., 2007).

Disease managers are faced with several challenges when responding to incursions of TADs. These include: what control measures to adopt; trade and economic implications of candidate control measures; how to manage resources such as personnel, equipment and vaccine; access to appropriate technology such as diagnostic tools; animal welfare issues; consumer concerns; and possible public health ramifications (Garner et al., 2007). The choice of control measures can be a compromise between the requirement for large-scale implementation and what is logistically, economically, and socially feasible (Tildesley et al., 2006; Mort et al., 2005). Epidemiological models are increasingly being employed as decision support tools for outbreak planning and response (Garner and Hamilton, 2011). Models are especially useful when a country has not recently experienced the disease of concern (Bates et al., 2003). The development and use of livestock disease spread models are quite often oriented to specific diseases in specific countries, for example, CSF in the Netherlands (Jalvingh et al., 1999), FMD in the Netherlands (Backer et al., 2012), FMD in Denmark (Boklund et al., 2013), ASF in Denmark (Halasa et al., 2018), PPR in Ethiopia (Fournié et al., 2018), HPAI in France (Andronico et al., 2019). This is understandable as funding for the development and use of decision support tools is usually provided in the context of a specific national interest. Difficulties can arise when extending models of the spread and control of livestock disease across national borders. Details on livestock population, production systems and marketing systems of neighbouring countries are not always readily available. It can also be a challenge to capture heterogeneities in production systems, control policies, and response resourcing across multiple countries, in a single transboundary model.

In 2017 the European Commission for the Control of Foot-and-Mouth Disease (EuFMD) commissioned a pilot study to develop a transboundary model of the spread and control of livestock disease in Europe, with FMD as the test case. The proposed model would be used to:

- study the size, duration, and economic impact of FMD outbreaks at both a single and multi-country scale
- assess the potential for establishment and spread of FMD under local conditions
- compare effectiveness and cost effectiveness of different candidate response strategies
- consider surveillance and early detection issues
- look at resource needs and resource management issues associated with managing outbreaks
- support training activities and simulation exercises

The model development project was undertaken with the support of the state veterinary authorities of seven central European member states of the EU: Austria, Bulgaria, Croatia, Hungary, Italy, Romania, and Slovenia. In this paper we describe a European livestock disease modelling framework EuFMDiS. EuFMDiS is a multi-country adaptation of the Australian Animal Disease Spread (AADIS) modelling framework (Bradhurst et al., 2015). AADIS is a national-scale disease modelling platform designed to provide decision support in the development of animal health policy in Australia. It captures livestock disease epidemiology, regional variability in transmission (e.g., due to environmental differences and seasonal livestock production and marketing patterns), and multi-jurisdictional approaches to control. EuFMDiS is a continental-scale modelling platform of livestock disease spread and control that simulates transmission within and between countries. It has been designed to support emergency animal disease planning in Europe. The outcome of the development project was the EuFMDiS modelling framework and the pilot EuFMDiS-FMD model.

## 2. MODEL OVERVIEW

### 2.1. Representation of a livestock population

The epidemiological unit in EuFMDiS is the ‘herd’, defined as a group of co-mingling animals of the same species under the same production system. A herd has static attributes such as herd type, species, location and jurisdiction, and dynamic attributes such as disease and vaccination states. A central assumption in EuFMDiS is that the livestock population in a study area can be categorised by ‘herd type’, such that key differences in production system characteristics and buying and selling patterns, can be satisfactorily captured. The stratification of the livestock population by herd type will largely be driven by the granularity of available data on livestock movements. It can be a challenge to define common herd types that apply across multiple countries where production systems and environments may vary considerably. EuFMDiS allows a user to define custom herd types appropriate to the disease being modelled and the study area. The set of herd types for the pilot EuFMDiS-FMD model (Table 1) was chosen through consultation with the participating countries. A EuFMDiS ‘holding’ is a collection of one or more co-located herds under the same management. This organisational structure allows EuFMDiS to represent the increased probability of disease transmission between herds that are co-resident on the same holding, due to the higher potential for direct contact and indirect contact via shared equipment and personnel.

**Table 1.**
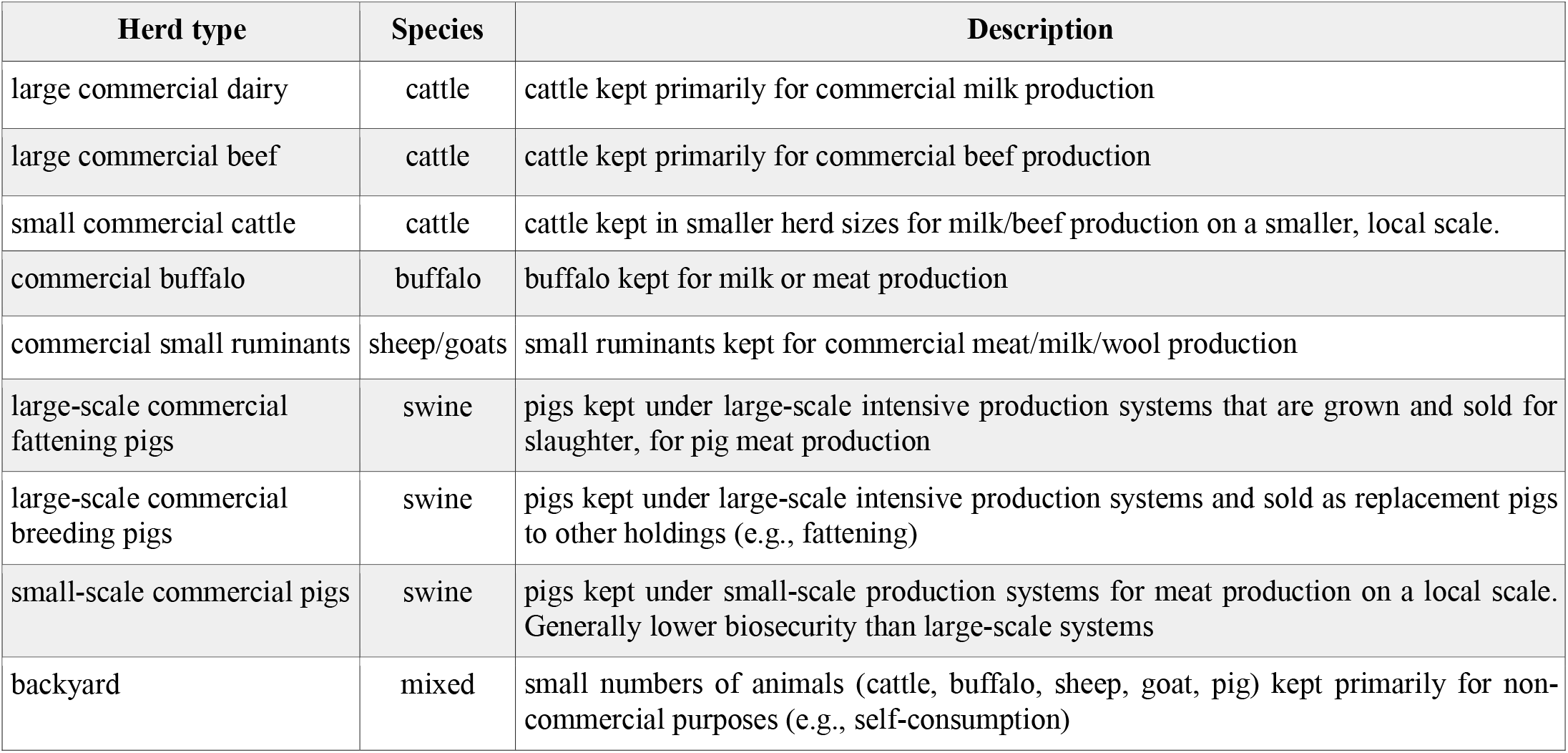
Herd types used in the pilot EuFMDiS-FMD model

Whilst EuFMDiS users have the option of employing a herd dataset comprising actual holdings at actual locations, this can give rise to privacy concerns (European Union, 2016). An alternative is to use a synthesized dataset based on census data or to obfuscate the identities of holding by perturbating their locations. Both techniques were employed during the assembly of herd data for the pilot EuFMDiS-FMD model.

### 2.2. Geospatial representation of a study area

A EuFMDiS study area is comprised of one or more countries. Each country is partitioned into one or more geographical ‘regions’ that are used to characterise regional heterogeneities in livestock production and marketing systems. For example, consider a country that is partitioned into three EuFMDiS regions: mountains, coastal, and plains. EuFMDiS allows a large commercial beef herd in the coastal region to have direct and indirect movements that are quite distinct from large commercial beef herds in the mountains and plains regions. 25 regions were defined for the EuFMDiS-FMD pilot model in consultation with the seven participating countries (Figure 1 and Table 2).

**Figure 1.**
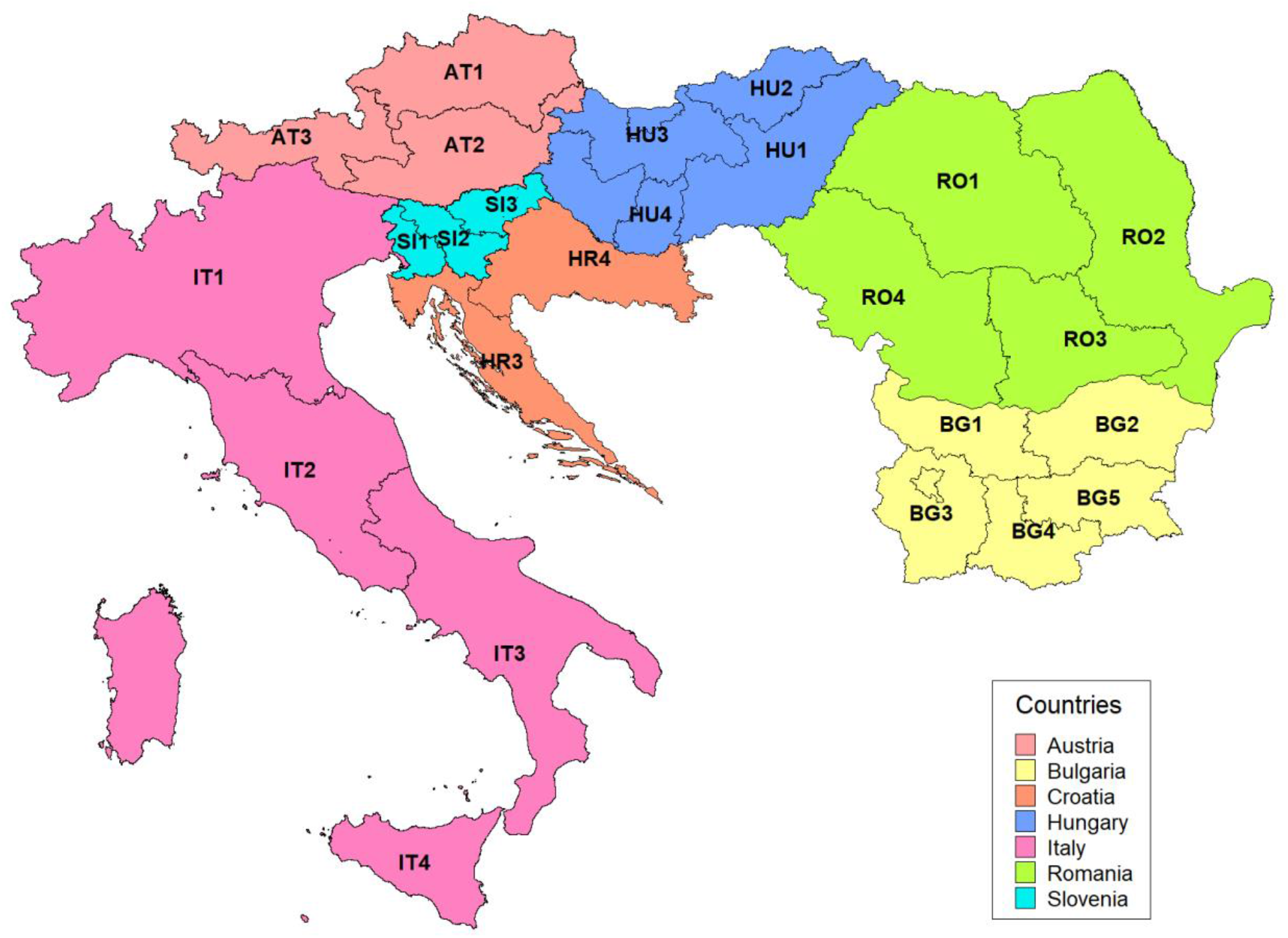
Regions defined for the pilot EuFMDiS-FMD model.

**Table 2.** Regions defined for the pilot EuFMDiS-FMD model

| Region | Country | Description |
| --- | --- | --- |
| AT1 | Austria | Austria north |
| AT2 | Austria | Austria south |
| AT3 | Austria | Austria west |
| BG1 | Bulgaria | Northwest Bulgaria |
| BG2 | Bulgaria | Northeast Bulgaria |
| BG3 | Bulgaria | Southwest Bulgaria |
| BG4 | Bulgaria | South Bulgaria |
| BG5 | Bulgaria | Southeast Bulgaria |
| HR3 | Croatia | South Croatia |
| HR4 | Croatia | North Croatia |
| HU1 | Hungary | Great Hungarian Plain |
| HU2 | Hungary | Hungary mountain areas |
| HU3 | Hungary | West-central Hungary |
| HU4 | Hungary | Transdanubian pig areas |
| IT1 | Italy | North Italy |
| IT2 | Italy | Central Italy |
| IT3 | Italy | South Italy |
| IT4 | Italy | Italian islands |
| RO1 | Romania | Romania N-V and Center |
| RO2 | Romania | Romania N-E and S-E |
| RO3 | Romania | Romania S and Bucharest |
| RO4 | Romania | Romania S-V |
| SI1 | Slovenia | Slovenia west |
| SI2 | Slovenia | Slovenia center |
| SI3 | Slovenia | Slovenia east |

### 2.3. Equation-based modelling of within-herd disease spread

EuFMDiS considers a herd to be homogeneous and ‘well-mixed’ from a disease transmission point of view, i.e., all members of a herd are deemed biologically equivalent and equally likely to contract a disease (Keeling and Rohani, 2008). EuFMDiS employs an SEIRDC (susceptible, exposed, infectious, recovered, deceased, clinical) compartmental equation-based model (EBM) to represent within-herd spread of the disease under study (Figure 2). The EBM can be thought of as comprising an SEIRD infection model (where exposed animals become infectious and then either recover or die), and a parallel SEC disease model (where exposed animals go on to either develop clinical disease or are asymptomatic). This approach is simple mathematically and is agnostic as to whether the latent period is less than the incubation period (i.e., there may be presymptomatic infectious animals), or whether the latent period is greater than or equal to the incubation period. Each infected herd has a system of ordinary differential equations (ODEs) (Equations 1–6), customised for the herd type and the pathogen of interest (Table 3).

**Figure 2.**
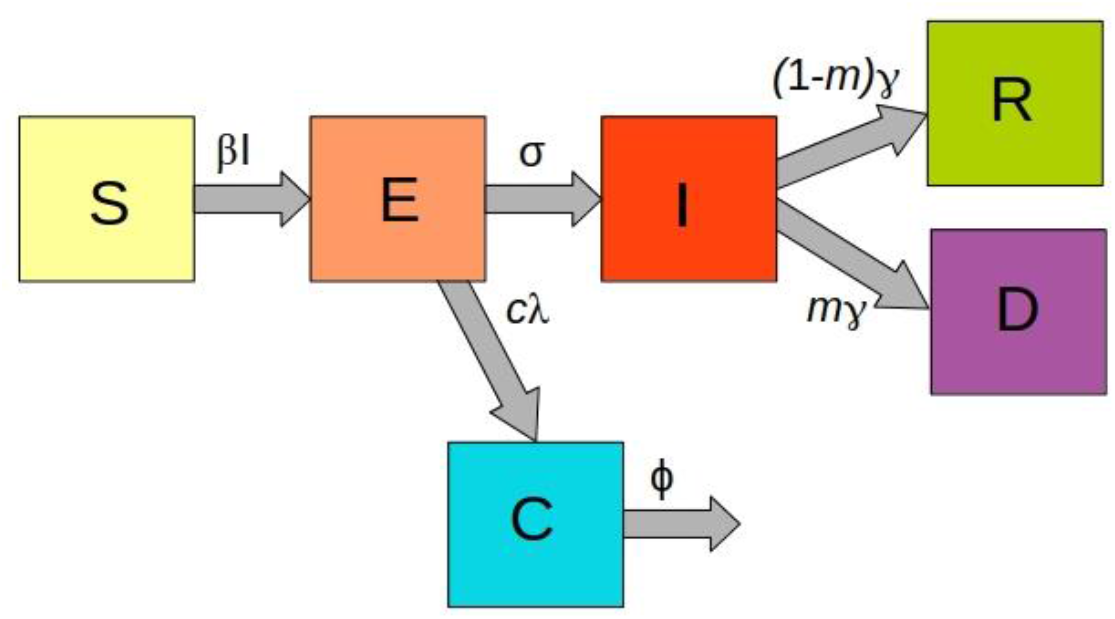
SEIRDC compartmental model used in EuFMDiS to represent the within-herd spread of disease.

**Table 3.** Parameterisation of the within-herd EBM used in the pilot EuFMDiS-FMD model

| Herd type | Tx rate ($\beta$) | Latent period (days) | Infectious period (days) | $R_0$ | Probability of mortality (m) | Incubation period (days) | Clinical period (days) | Proportion clinical (c) |
| --- | --- | --- | --- | --- | --- | --- | --- | --- |
| large commercial dairy | 2.1 | 2 | 10 | 21.0 | 0.05 | 3 | 12 | 1 |
| large commercial beef | 1.6 | 2 | 10 | 16.0 | 0.05 | 3 | 12 | 1 |
| small commercial beef | 1.8 | 2 | 10 | 18.0 | 0.05 | 3 | 12 | 1 |
| commercial buffalo | 2.0 | 2 | 10 | 20.0 | 0.02 | 4 | 12 | 1 |
| commercial small ruminants | 0.25 | 5 | 10 | 2.5 | 0.03 | 6 | 10 | 0.5 |
| large-scale commercial fattening pigs | 2.2 | 1 | 6 | 13.2 | 0.03 | 5 | 14 | 1 |
| large-scale commercial breeding pigs | 2.2 | 1 | 6 | 13.2 | 0.15 | 5 | 14 | 1 |
| small-scale commercial pigs | 2.1 | 1 | 6 | 12.6 | 0.1 | 5 | 14 | 1 |
| backyard | 1.5 | 4 | 5 | 7.5 | 0.03 | 4 | 12 | 0.75 |

EuFMDiS simplifies herd size by assuming that inflows (births and transfers in) are equivalent to outflows (non-disease related deaths and transfers out). When a susceptible herd becomes infected the ODE system is solved numerically to yield the SEIRDC compartmental counts over time. The EBM generates curves predicting the infected, infectious, and clinical prevalence of the infected herd. The solution remains in place up until an external event (such as vaccination or culling), acts upon the EBM. If a herd is vaccinated and immunity levels increase, the EBM reacts by resolving the ODE system to yield updated SEIRDC compartment counts from that point in time onward.

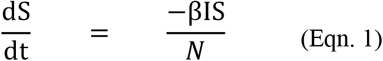

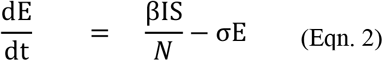

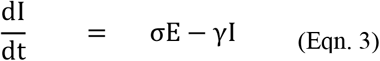

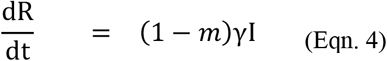

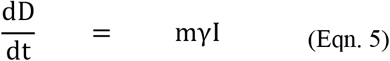

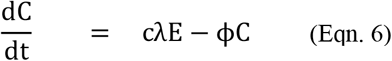

where S = number of animals in the herd that are susceptible

E = number of latently infected animals in the herd

I = number of infectious animals in the herd

R = number of recovered animals in the herd

N = total number of animals in the herd (= S + E + I + R)

D = number of animals in the herd that have died from the disease

C = number of animals in the herd with clinical signs

β = effective contact rate (contact rate × transmission probability)

σ = infectious progression rate (1/σ = average duration of the latent period)

γ = recovery rate (1/γ = average duration of the infectious period)

m = average probability of disease-related mortality c = proportion of cases showing clinical signs

λ = clinical disease rate (1/λ = average duration of the incubation period)

φ = clinical recovery rate (1/φ = average duration of the clinical period)

### 2.4. Agent-based modelling of between-herd disease spread

EuFMDiS represents the spread of disease between herds with a stochastic and spatially-explicit agent-based model (ABM) (Bradhurst, 2015). The levels of infected and infectious prevalence predicted by a herd’s EBM inform the likelihood that between-herd spread will occur. EuFMDiS provides two distinct options for spreading disease between herds:

1. <u>Data-driven spread pathways</u> - there are six independent data-driven pathways involving direct contacts (animal movements) or indirect contacts: (i) local spread (ii) direct spread between holdings (iii) direct spread via markets (iv) transboundary direct spread via assembly centres (v) indirect spread and (vi) airborne spread. An additional pathway for seasonally shared pastures is under development. These spread pathways require detailed parameterisation and are dependent on the availability and quality of the underlying data.
2. <u>Analytical spread pathways</u> - a diffusion pathway represents localised spread of infection around an infected holding while a jump pathway simulates longer distance jumps of infection to new locations. These pathways are much simpler to parameterise and can be useful when there is inadequate data to support the data-driven spread pathways.

The choice between data-driven and analytical spread pathways is made on a per-country basis. This means it is possible for disease transmission to switch between the data-driven and analytical approaches at national borders.

Each spread pathway has a stochastic algorithm that determines on any given simulation day whether disease transfers from infectious herds to susceptible herds (Bradhurst, 2015). The EuFMDiS spread pathways are described below in the context of EuFMDiS-FMD.

#### Local spread

Local spread is a catch-all pathway for very short-range transmission of disease from an infected herd to nearby susceptible herds (Sanson, 1994). Local spread can be an important pathway in high-density farming areas, for example, most of the cases in the 2001 UK FMD outbreak were attributed to local spread (Gibbens et al., 2001). The underlying mechanisms of local transmission may include: short-range aerosol spread across fences; direct spread via the straying of stock; and indirect spread via vehicles, people, surface runoff, and sharing of equipment between neighbours (Gibbens et al., 2001; Kitching et al., 2006).

EuFMDiS represents local spread with a spatial kernel that aggregates all spread mechanisms inside a circular area enclosing each infected herd (default radius 3km). To avoid double-counting, the other data-driven spread pathways do not operate inside the local spread area. All susceptible herds inside a local spread area are deemed at-risk. The probability of transmission from an infected ‘source’ herd to each at-risk ‘destination’ herd is decided stochastically, taking into account: infectious prevalence in the source herd; infectivity of the source herd (based on species and size); susceptibility of the destination herd; biosecurity measures in place at the destination holding; and the distance between the source and destination herd (Equation 7).

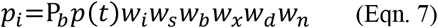

where *p_i_* = probability that a local contact between herds results in infection

*P_b_* = baseline probability that a local contact between herds results in infection (per region)

*p(t)* = normalised infectious prevalence of the source herd at time *t*

*w_i_* = infectivity weight of the source herd

*w_s_* = susceptibility weight of the destination herd

*w_b_* = biosecurity weight of the destination herd (user configurable constant reflecting the influence of varying biosecurity practices across herd types on the risk of exposure to disease)

*w_x_* = seasonal weight (user configurable constant reflecting how varying environmental conditions influence virus survival, defined per month, per region)

*w_d_* = distance weight (reflecting the influence of proximity to an infection source on the risk of exposure)

*w_n_* = detection weight (user configurable constant reflecting that local spread may organically dampen once an outbreak has been declared due to an increased awareness of risk of transmission leading to decreased movements of people and vehicles, etc.)

The distance weight *w_d_* can be configured to decay linearly (Equation 8) or exponentially (Equation 9).

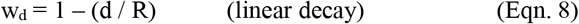

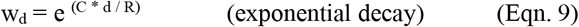

where
d = distance from the source herd to the destination herd

R = diffusion radius (user configurable, default 3 km)

C = decay constant (user configurable, default −3.4539)

Local spread can also occur between herds that are co-resident on the same holding. In this case the baseline probability of transmission *P_b_* is increased to reflect the higher potential for direct and indirect contacts between herds managed on the same holding.

Tildesley and colleagues (2012), found that a non-linear relationship between herd size and infectivity/susceptibility better described data from the 2001 UK FMD outbreak than a linear relationship. EuFMDiS provides user-configurable power law parameters *P_i_* and *P_s_* that specify the level of influence that herd size has on infectivity and susceptibility. Infectivity weights depend on species and herd size and are scaled across the herd population (Equation 10). The relative infectivity constants *Si* specify the infectivity of a species to FMD, relative to sheep (Risk Solutions, 2005). The infectivity powers *P_i_* allow per-species tuning of the effect of herd size on infectivity (0 ≤ *P_i_* ≤ 1, where a value of 0 specifies no effect and a value of 1 specifies a linear relationship).

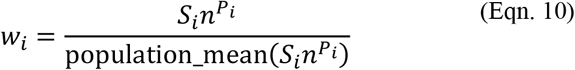

where *w_i_* = infectivity weight

*S_i_* = species relative infectivity (user configurable, defaults: sheep=1, cattle=2, pigs=4)

*n* = herd size

*P_i_* = species infectivity power (user configurable, defaults: sheep=0.35, cattle=0.35, pigs=0.35) Susceptibility weights also depend on species and herd size and are scaled across the herd population (Equation 11). The relative susceptibility constants *S_s_* indicate the susceptibility of a species to FMD, relative to sheep (Risk Solutions, 2005). The susceptibility powers *P_s_* allow per-species tuning of the effect of herd size on susceptibility (0 ≤ *P_s_* ≤ 1, where a value of 0 specifies no effect and a value of 1 specifies a linear relationship).

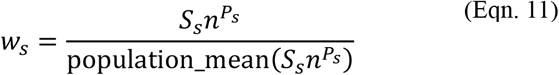

where
*w_s_* = susceptibility weight

*S_s_* = species relative susceptibility (user configurable, defaults: sheep=1, cattle=6, pigs=0.4)

*n* = herd size

*P_s_* = susceptibility power (user configurable, defaults: sheep=0.35, cattle=0.35, pigs=0.35)

When a susceptible herd becomes infected, an EBM is created and solved with initial conditions based on the estimated number of exposed animals in the destination herd and the size of the destination herd.

#### Direct contact spread

TADs can spread when an infectious animal comes into direct contact with a susceptible animal. For example, respiratory transmission can occur between animals sharing a paddock, yard, pen or truck. Direct contact spread through the relocation of live animals is often reported as the most significant means of transmission of TADs (Gibbens et al., 2001; Kao, 2002; Green et al., 2006; Kao et al., 2006; Kao et al., 2007; Lindström et al., 2009; Kitching, 2011). EuFMDiS simulates animal movements from infected herds. The daily likelihood of a consignment leaving an infected herd is derived from livestock movement frequency data that depends on herd type, region, and season. The destination of the consignment (another holding, a slaughterhouse, a market, or export), is determined stochastically from livestock movement data that depends on the source herd type and region.

##### (i) Movements between holdings

The destination region, destination herd type, movement distance and destination herd are determined stochastically from livestock movement data that depends on the source herd type and region. Transmission depends on the prevalence of infection in the source herd and the consignment size (Equation 12).

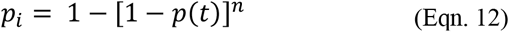

where
*p_i_* = probability that a consignment contains at least one exposed or infectious animal

*p(t)* = infected prevalence in the source herd at time *t* (as modelled by the EBM)

*n* = consignment size

When a susceptible herd becomes infected an EBM is created and solved with initial conditions based on the proportion of infectious and exposed animals in the consignment, and the size of the destination herd.

##### (ii) Movements from holdings to slaughterhouses

Movements from infected holdings to slaughterhouses are logged but no further spread occurs, i.e., they are considered ‘dead-ends’ with respect to disease transmission.

##### (iii) Movements from holdings to markets

Markets have the potential to greatly amplify an outbreak prior to the disease being recognised and controls implemented (Gibbens et al., 2001). The transmission of disease is facilitated by the stresses of transit and handling, large numbers of susceptible animals, and the mixing and partitioning of stock into consignments. Further, outgoing consignments can potentially carry infection to multiple widely dispersed locations. For example, the rapid escalation of the 2001 UK FMD outbreak was attributed to the unwitting movement of infected sheep to and from markets (Ferguson et al., 2001; Mansley et al., 2003; Mansley et al., 2011). A modelling study by Green and colleagues (2006) suggests that the number of infected markets strongly influences the eventual outbreak size.

Each EuFMDiS herd is assigned a market based on proximity to the herd, the type of herd, and the species typically processed at the market. At a market, animals from different sources may be mixed and sorted such that a single infected consignment entering a market may contribute to multiple infected consignments leaving the market. The destination of each infected consignment leaving the market (another holding, a slaughterhouse or export) is determined probabilistically from livestock movement data that depends on the source herd type and region. For consignments destined for other holdings, buyers are selected based on radial catchment areas. Infection is transmitted from infected consignments to destination herds with a force relative to the viral load in the consignment.

##### (iv) Transboundary movements via assembly centres

Assembly centres are places where animals from different holdings are held and prepared for onward consignment, predominantly to export markets. The following assumptions simplify modelling the transboundary movements of livestock:

- all consignments leaving a holding and destined for export are sent to an assembly centre.
- export consignments have specific destinations and do not undergo splitting and mixing whilst at the assembly centre.
- export consignments may be dispatched to a country within the study area or a country outside the study area. Destination countries outside the study area can be further distinguished as associated in some way (e.g., another EU member state) or not associated (e.g., a non-EU country). The destination country is chosen probabilistically based on the possible destinations for the consignment species and the country from which the consignment originated.
- consignments arriving in countries within the study area are held in assembly centres before being consigned onward to holdings or slaughterhouses, based on region and species. Destination holdings for consignments are chosen stochastically based on region and source herd type. Consignments sent to slaughterhouse are logged but no further spread occurs.
- consignments arriving in countries outside the study area are logged but no further spread occurs.

Figure 3 provides an overview of the transboundary direct spread pathway for the pilot EuFMDiS-FMD model. The livestock movement data informing within-country direct spread were provided by the participating countries from national databases. The livestock movement data informing transboundary spread were sourced from the Trade Control and Expert System (TRACES) system (European Commission, 2020). TRACES is an online management tool for all sanitary requirements on intra-EU trade and importation of animals, feed, semen and embryo, food, and plants.

**Figure 3.**
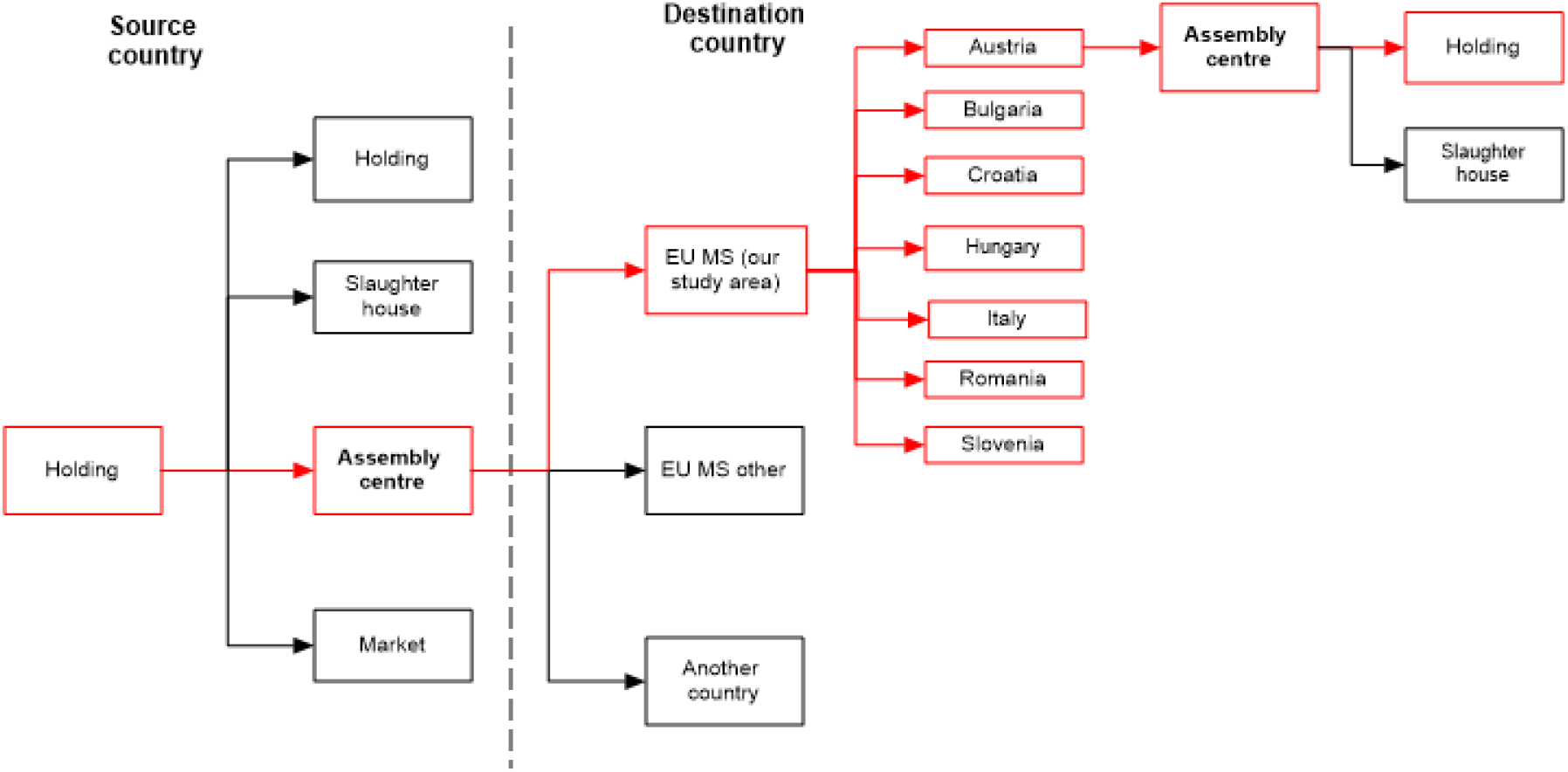
EuFMDiS-FMD transboundary direct spread pathway.

#### Indirect contact spread

Indirect contact transmission arises from the movement between herds of contaminated animal products, by-products, and fomites such as equipment, people and vehicles. Potential sources include veterinarians, shearing contractors, artificial insemination technicians, milk tankers, and stock feed delivery vehicles. Indirect contacts can be categorised as high, medium or low according to their potential for transmitting infection (Nielen et al., 1996; Bates et al., 2001; Sanson, 2005; Nöremark et al., 2013). EuFMDiS employs a single category of indirect contacts with a specified average (baseline) probability of transmission. The user can parameterise indirect spread to represent different risk profiles. Compared to direct contacts, there is limited data on indirect contacts. The type and location of exposed herds is determined stochastically using a contact matrix and distance distributions by herd type. If a herd is exposed through indirect contact, the probability of transmission depends on the infectious prevalence of the source herd, the relative infectiousness of the source herd (based on species and herd size), environmental conditions that influence virus survival, biosecurity practices, and relative susceptibility of the exposed herd (based on species and herd size) (Equation 13).

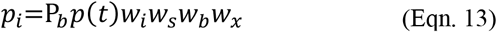

where
*p_i_* = probability that the indirect contact results in an infection

*P_b_* = baseline probability that an indirect contact results in infection

*p(t)* = normalised infectious prevalence of the source herd at time *t*

*w_i_* = infectivity weight of the source herd (per local spread)

*w_s_* = susceptibility weight of the destination herd (per local spread)

*w_b_* = biosecurity weight of the destination herd (per local spread)

*w_x_* = seasonal weight (per local spread)

#### Airborne spread

Airborne spread is the infection of susceptible animals by virus conveyed on the wind. In the case of FMD, pigs pose the greatest threat for airborne spread because of their potential to excrete large quantities of virus relative to other species (Donaldson and Alexandersen, 2002; Alexandersen et al., 2003). The extent of a viral plume depends on the concentration of virus in the source herd, weather conditions and the strain of virus (Donaldson et al., 2001; Donaldson and Alexandersen, 2002; Gloster et al., 2006). EuFMDiS adopts a simplified approach to airborne spread similar as per the AusSpread (Garner et al., 2006) and AADIS models (Bradhurst et al. 2015), with probabilities of the likelihood of airborne spread defined per weather station, per month. In the case of FMD, the most favourable meteorological conditions for airborne spread are constant wind direction, wind speed of 5 metres/second, high atmospheric stability, no precipitation, and relative humidity greater than 55% (Donaldson et al., 2001).

Locations of weather stations and weather data were sourced from the European Climate Assessment and Dataset (ECAD, 2019). During initial model setup for the disease under study, the EuFMDiS user specifies the set of herd types that may be capable of transmitting disease by airborne spread beyond the local spread area. In the case of FMD, this is limited to commercial pig herds. For each simulation day, the weather station closest to each candidate infectious herd is queried as to whether conditions are suitable for airborne spread. For each herd that is deemed a potential source of airborne spread, a sector is constructed in the prevailing wind direction for the month, subtended by a configurable angle of default size 30° (i.e., θ=15° on either side of the wind direction vector). The extent of a plume depends on the number of infectious animals in the source herd (Donaldson et al., 2001), and relates the number of virus-shedding animals to the distance downwind to susceptible animals (Equation 14). Topographical features such as mountains, lakes and forests that might influence a plume are not considered. In the case of FMD, although there have been reports of viral plumes travelling substantial distances in ideal conditions over open water (Donaldson et al., 1982), the anticipated maximum extent of a plume over land is in the vicinity of 10 to 20 km (Mikkelsen et al., 2003; Gloster et al., 2006; Schley et al., 2009).

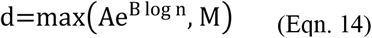

where *d* = distance of the viral plume

*n* = the number of infectious pigs in the source herd

*A* = plume coefficient (user configurable with default 0.113 for FMD)

*B* = plume exponent (user configurable with default 1.367 for FMD)

*M* = maximum distance of a viral plume (user configurable with default 20 km for FMD)

The probability of transmission to susceptible herds in the airborne spread sector takes into account the susceptible herd species, the size of the susceptible herd, and the distance of the susceptible herd from the infected herd (Equation 15) (Donaldson et al., 2001; Garner et al., 2006).

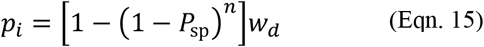

where *p_i_* = probability that a susceptible herd will become infected

*P_sp_* = probability that a single animal of a specific susceptible species will become infected

*n* = size of the susceptible herd

*w_d_* = distance weight (per local spread)

The distance weight *w_d_* represents the diffusion of a plume with distance from a source herd, and hence the diminishing risk of transmission. As per local spread, distance weight is configurable as having either linear or exponential decay.

In the case of EuFMDiS-FMD, the airborne spread pathway is disabled by default, but can be easily enabled if required.

#### Diffusion spread

The probability of transmission via diffusion spread is defined via a spatial kernel that can operate in three modes: linear, exponential and power-law (Equation 16). The linear and exponential diffusion modes operate in the same manner as the local spread pathway. Power-law diffusion is based on Backer et al., 2012 (Equation 17).

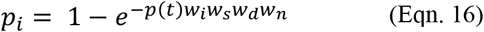

where *p_i_* = probability that the diffusion contact results in an infection

*p(t)* = normalised infectious prevalence of the source herd at time *t*

*w_i_* = infectivity weight of the source herd

*w_s_* = susceptibility weight of the destination herd

*w_d_* = transmission rate

*w_n_* = detection weight (per local spread)

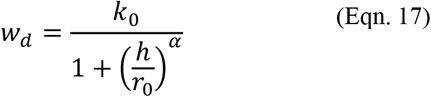

where *w_d_* = transmission rate

*h* = distance from the source herd to the destination herd

*k_0_* = transmission rate when *h* is zero

*r_0_* = distance at which the transmission rate is 0.5 k_0_

*α* = kernel shape parameter

#### Jump spread

The probability of a longer distance jump transmission (Equation 18) depends on the infectious prevalence of the source herd, the relative infectiousness of the source herd (based on species and herd size), environmental conditions that influence virus survival, biosecurity practices, and relative susceptibility of the exposed herd (based on species and herd size). The frequency with which longer distance jumps occur is set by the user.

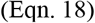

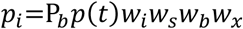

where *p_i_* = probability that the jump contact results in an infection

*P_b_* = baseline probability that a jump contact results in infection

*p(t)* = normalised infectious prevalence of the source herd at time *t*

*w_i_* = infectivity weight of the source herd (per local spread)

*w_s_* = susceptibility weight of the destination herd (per local spread)

*w_b_* = biosecurity weight of the destination herd (per local spread)

*w_x_* = seasonal weight (per local spread)

### 2.5. Disease control

The EuFMDiS unit of interest for disease control is the holding. A holding has static attributes such as holding type and the set of constituent herds, and dynamic attributes describing the status of control operations. The main simulated control strategies are movement restrictions, surveillance, tracing, infected holding operations (culling, disposal, and cleaning and disinfection), pre-emptive culling and emergency vaccination. Control measures are defined and resourced per country.

EuFMDiS control measures are configurable for the disease of interest. They are described below in the context of EuFMDiS-FMD with configuration that is consistent with the European FMD Directive (European Union, 2003).

#### Detection of the index case

The control and eradication phase of an outbreak commences after the declaration of the index case, i.e., the first declared infected holding (IH). The day of first detection is either determined stochastically (using pre-configured probabilities of reporting by herd type, and clinical prevalence), or occurs on a fixed day at a specific or randomly selected holding.

#### Movement Restrictions

Following the detection of the first IH, EuFMDiS provides the option of a national livestock standstill, where direct movements are halted for a configurable number of days in the affected country. A user-defined compliance percentage allows for the possibility of illegal movements occurring during the standstill. Controlled areas are established around each IH in order to restrict the movement of livestock, products and other material. The controlled areas are defined and enforced per country, and may be designated areas (local administrative area, entire country), or radius-based per IH. There are two levels of control: Protection Zones (PZs) that immediately enclose IHs, and Surveillance Zones (SZs) that enclose PZs. PZs have the highest level of control while SZs have a lower level of control. The default settings are 3 km and 10 km radii around IHs for PZ and SZ respectively, per the EU FMD Directive (European Union, 2003). Radial controlled areas are clipped to fall within the national boundaries of the subject IH. When IHs are clustered a meta-PZ and meta-SZ are formed from the union of the constituent PZs and SZs.

#### Surveillance

Surveillance is the process by which new infections are identified and declared. During a TAD outbreak, surveillance is used to detect new outbreaks, define the extent of infection, and demonstrate freedom in uninfected areas. EuFMDiS allows for reporting of suspect cases on an ad hoc basis by owners/inspectors or others. This represents one of the most important mechanisms for finding new infected holdings (McLaws et al., 2007). EuFMDiS commences suspect case reporting the day after the first IH has been declared and allows for both true positive and false positive reports. False positive reports identify herds that are exhibiting symptoms but are not actually infected with the subject disease. True positive reports are generated stochastically based on an infected herd’s clinical prevalence and the probability of reporting (which depends on herd type). The EuFMDiS user configures whether surveillance visits will include a laboratory test or not, and if so, the delay in days before a result will be available. The number of false positive reports generated is proportional to an n-day (default n=3), moving average number of true positive reports. The modelling of both true and false reports facilitates more realistic modelling of surveillance as resources are consumed regardless of whether a surveillance visit yields a positive assessment or not. EuFMDiS also models the active inspection of at-risk holdings within a designated distance of IHs, subject to a configurable inspection schedule (number and frequency of inspections).

#### Tracing

Tracing is the identification of movements onto and off IHs in order to ascertain where infection may have come from or gone to. Tracing includes animals, products, equipment, vehicles and people. Traced holdings may be true cases (and thus infected), or false (not infected). EuFMDiS identifies true traces by following infection chains during a simulation, allowing for variable tracing effectiveness by species and pathway (direct contact versus indirect contact), and tracing duration. False forward traces are obtained by applying the direct and indirect spread pathways to a holding of interest within the forward tracing window. False backward traces are obtained by reversing the direct and indirect spread pathways over the backwards tracing window (i.e., modelling movements onto holdings of interest). This approach results in a set of plausible false traces, i.e., holdings of a suitable type and location that could well have been sources or destinations of movements of concern.

Holdings that require visits by surveillance teams are identified through tracing, active inspection of holdings within PZs and reporting of suspect holdings. EuFMDiS maintains a dynamic queue of holdings awaiting a surveillance visit. Surveillance visits are prioritised according to a configurable scheme that considers holding classification, declared area and herd type. If multiple holdings have the same priority, then arbitration is based on how long a holding has been waiting for a visit. The visit duration (based on herd type), visit frequency (based on priority), and overall surveillance period are configurable.

#### IH operations

IH operations are the valuation, destruction and disposal of animals (‘stamping out’) and decontamination. Stamping out is the default policy for controlling an outbreak of FMD as it is considered the fastest way to reduce viral excretions and dampen spread. All IH operations are prioritised based on holding type and herd size. The times required for a holding to undergo culling, disposal and decontamination are defined by herd type in the EuFMDiS configuration data.

#### Pre-emptive culling

EuFMDiS also provides the option of ring culling holdings within a configurable distance of each IH, and the pre-emptive culling of holdings that are deemed high risk because of a traced direct contact with an IH. Pre-emptive culling operations are prioritised based on the reason for culling (stamping out takes precedence over pre-emptive culling), holding type, herd size, and distance to the nearest IH.

#### Emergency vaccination

Emergency vaccination strategies include:

- <u>Suppressive</u> – vaccination is carried out inside known infected areas in order to suppress virus production in at-risk and exposed herds and dampen further spread.
- <u>Protective</u> – vaccination is carried out outside known infected areas in order to protect susceptible animals from infection.
- <u>Mass</u> – vaccination is carried out across a broad area to large numbers of animals. This strategy could be applied if an outbreak is not under control and there is a risk of spread escalating.

EuFMDiS provides several triggers for commencing a vaccination program: i) on a configurable day into the control program; ii) once a configurable number of IHs has been declared; iii) once a pending cull threshold has been reached. EuFMDiS models all vaccination policies with an annulus of configurable inner and outer radii. The inner radius is set to zero for suppressive and mass vaccination. A vaccination annulus is established around each target IH, and eligible holdings inside the annulus are scheduled for vaccination. The user can select to only vaccinate around IHs found on or after the day the vaccination program begins, or around all new and previously identified IHs. The vaccination candidates inside each annulus are prioritised according to herd type, herd size, and proximity to the nearest IH. It is also possible to omit certain herd types from vaccination. The direction of vaccination (from the outside in, or from the inside out), is set in the EuFMDiS configuration data. EuFMDiS also includes the option of selectively vaccinating herds in high-risk areas only. This involves pre-defining high-risk areas and flagging herds within these areas.

The effect of vaccination is to increase immunity in a herd over time. When a partially immune herd is exposed to infection, the virus production profile generated by the EBM reflects that some of the animals have protective immunity. The level of protection will depend on the timing of the vaccination program (in relation to infection exposure events), the configured vaccine efficacy, and the configured vaccine immunity profile that governs the waxing and waning of immunity over time. Vaccination visits to holdings are prioritised according to herd type, herd size and proximity to an IH. The time required for a holding to undergo vaccination is defined by herd type in the EuFMDiS configuration data.

#### Post-outbreak management

Disease models often stop simulating once an outbreak has been controlled i.e., all infected herds have been found and the control program has concluded. However, from a disease manager’s perspective additional work is required before a country can regain disease-free status. This will include decisions about managing vaccinated animals in the population and undertaking the surveillance necessary to support disease freedom. EuFMDiS provides three policy options for the post-outbreak management of vaccinated animals:

- retention in the population to live out normal commercial lives
- removal from the population for disposal to waste
- removal from the population for salvage

The choice of policy influences post-outbreak resource requirements and compensation payments (if vaccinated animals are removed), and the length of the mandatory OIE waiting period before return to trade (Bradhurst et al., 2019).

Post-outbreak surveillance is conducted in terms of ‘clusters’ that represent discrete areas of previously declared infection. A cluster is formed from the union of overlapping SZs. For example, Figure 4 illustrates five clusters, where the black dots are previously infected holdings, the red areas are the grouped PZs and the green areas are the grouped SZs. Post-outbreak surveillance is carried out independently in each cluster in order to provide support for proof-of-freedom. A user-defined sampling regime determines the number of herds to test within a cluster, and the number of animals to test within a selected herd, in order to achieve statistical confidence that residual infection would be detected. For example, a 95:5 sampling regime implies that sufficient herds are randomly tested in a cluster in order to achieve 95% confidence that a residual infected prevalence of at least 5% would be detected (Cannon and Roe, 1982; Bradhurst et al., 2021b).

**Figure 4.**
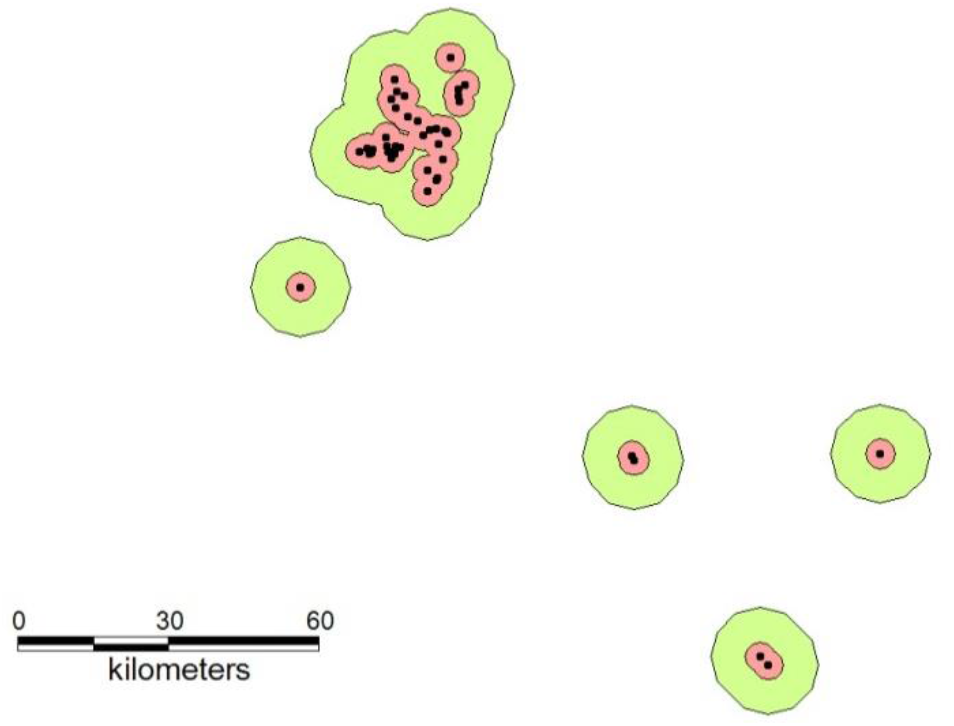
Example of infection clusters in which EuFMDiS post-outbreak surveillance is carried out.

Testing regimes are defined in terms of screening and confirmatory tests that depend on herd type and whether vaccination was used during the control program (i.e., whether structural protein tests or non-structural protein tests are appropriate). Tests may be a clinical, serological, or virological, and are defined in terms of sensitivity, specificity, cost, throughput, and pooling rate. The latter allows for the incorporation of new approaches and pooled tests such as bulk milk testing and pig salivary ropes (Armson et al., 2018; Grau et al., 2015). EuFMDiS reports the number of true/false positives and true/false negatives, and the duration and cost of the post-outbreak surveillance program.

It can be very challenging for a disease manager to decide when the final IH of an outbreak has been declared and processed, and post-outbreak activities should commence. EuFMDiS represents this with a user-defined rolling countdown timer (e.g., 30 days) that starts whenever a new IH is declared and processed. If the countdown timer expires then the outbreak is assumed over, and post-outbreak management and surveillance activities commence.

#### Resourcing

The resources required to manage an emergency animal disease outbreak include personnel (e.g., veterinarians, animal health officers, control centre staff), equipment (e.g., vehicles), facilities (e.g., laboratories) and consumables (e.g., vaccine, disinfectant). Some aspects of disease control and eradication are resource intensive, and the lack of resources can severely hamper the response to an outbreak (Roche et al., 2014). EuFMDiS models the resources required for key operational activities: surveillance, culling, disposal, decontamination, and vaccination. An EuFMDiS ‘resource’ is abstract in that it can represent whatever is required to complete a specific task. For example, the resource required to conduct a surveillance visit might be a veterinarian, an assistant, and a vehicle. As countries are responsible for emergency animal disease management within their own boundaries, resources are organised into pools by jurisdiction, (i.e., each country has five pools covering the key operational activities).

When a field operation is scheduled, a resource is requested from the relevant pool of the country. If a resource is available, then it is ‘borrowed’ from the pool and the field operation commences. If a resource is not available, then the field operation is queued until such time as a resource becomes available. Once a field operation has completed, the resource is ‘returned’ to the pool. It is anticipated that the resources available to manage a disease outbreak ramp up over time, so initially the pools are small and increase in a linear manner up to a maximum size. The starting point, duration of the ramp-up and maximum pool size are defined in the EuFMDiS configuration data, by resource type and by country. EuFMDiS tracks the availability and allocation of resources to provide immediate feedback as to whether/where the control program is resource constrained. Resource pools can be configured to be ‘unlimited’ in which case resources are always immediately granted upon request. In this mode the resourcing profile of an outbreak is a model output that conveys the level of resourcing required, rather than a constraint on the efficacy of the control program.

#### Outbreak costs

While detailed economic evaluations and optimisations are generally outside the scope of epidemiological models, EuFMDiS provides some useful costing outputs. These provide insight into the potential economic impact of an outbreak and allow relative cost-benefit comparisons of different control strategies. EuDMDiS keeps track of control costs (control centre operations, field operations, compensation, vaccine), post-outbreak management costs (control centre operations, field operations, compensation), and potential loss of trade. The latter is estimated simply from the value of a country’s exports of animals and animal products, and the number of days from the declaration of the index case through to the end of the mandatory OIE waiting period.

### 2.6. Implementation

The EuFMDiS and AADIS modelling frameworks utilise a common agent-based modelling platform (Bradhurst, 2015) which can operate in four modes: contagious livestock disease, vector-borne livestock disease, plant/environmental pests, and human disease. When modelling the spread and control of contagious disease in livestock, the agents are herds, holdings (comprising one or more herds), markets and slaughterhouses. When modelling the spread and control of plant and environmental pests, the agents are cells in a lattice environment (Bradhurst et al., 2021a). When modelling the spread and control of insect vector-borne livestock disease (such as bluetongue), the agents are herds, holdings, markets, slaughterhouses *and* cells. When modelling the spread and control of human disease the agents are people. Descriptions of the vector-borne livestock and human disease modes will appear in future papers.

EuFMDiS agents are lightweight and threadless which scales well with livestock population size. The agents interact in a spatially-explicit environment comprised of disease spread and control components that operate concurrently and independently (Bradhurst, 2015). As the environment components are independent and operate on dedicated threads, they can all be separately enabled/disabled. This allows EuFMDiS to easily switch between disaggregated data-driven spread modelling and aggregated analytical spread modelling. The implementation of each component is private and alternate components can be swapped in and out. For example, the implementation of the surveillance component can change without triggering a cascade of software modifications to other model components.

EuFMDiS is written in Java (Oracle, 2020), and employs open-source products including SQL Power Architect (SQL Power Group, 2020), PostgreSQL (PostgreSQL, 2020), OpenMap (BBN, 2016) and Log4j (Apache, 2020). EuFMDiS runs under either Linux™ or Windows™ and has an asynchronous software architecture with concurrency achieved through Java threads. The ability to take advantage of cheap parallelism available with multi-core personal computers along with a hybrid model architecture, in-memory database, and grid-based spatial indexing, allow EuFMDiS to efficiently conduct large-scale simulations (Bradhurst et al., 2016).

The primary EuFMDiS outputs are comma-separated value files, which can be post-processed statistically. EuFMDiS also provides a graphical user interface for interacting with the model and dynamic visualisation of incursions as they unfold (Figure 7). The ability for EuFMDiS to convey incursion and management concepts visually has proven effective in a classroom setting. The ability of the model to contrast the full extent of a simulated outbreak with the limited view that a disease manager has may also be useful for communication with decision makers and outbreak simulation exercises.

### 2.7. Verification and validation

As the EuFMDiS and AADIS modelling frameworks have a common underlying code baseline, EuFMDiS inherits from previous AADIS verification and validation activities, and modelling studies (Bradhurst, 2015; Bradhurst et al., 2015; Bradhurst et al., 2016; Garner et al., 2016; Van Andel et al., 2018; Bradhurst et al., 2019; Firestone et al., 2019; Firestone et al., 2020).

An independent review of the EuFMDiS model was conducted by Wageningen Bioveterinary Research to assess fitness for purpose for modelling in a European context (de Vos et al., 2019). EuFMDiS was described as a great help in contingency planning by allowing evaluation of different control strategies, ranging from the minimum requirements set by EU legislation up to the inclusion of ring culling and/or vaccination, while also considering limited availability of resources. The review made many helpful recommendations including the need for better documentation of input parameters and outcome variables, and the need for further model validation and sensitivity analysis of input parameters.

Validation of the EuFMDiS model will be an ongoing process through various case studies and sensitivity analyses being conducted by the EuFMD, participating EU member states, and post-graduate students (for example, Marschik et al., 2021).

## 3. CASE STUDY

The following simple epidemiological case study is provided to illustrate how EuFMDiS-FMD can be used to address policy issues.

FMD is an acute, highly contagious viral disease of domestic and wild cloven-hoofed animals. The disease is clinically characterised by the formation of vesicles and erosions in the mouth and nostrils, on the teats, and on the skin between and above the hoofs (Meyer and Knudsen, 2001). The FMD virus spreads between hosts through direct contact (e.g., movement of live animals between holdings, and between holdings and markets), indirect contact (e.g., livestock products, byproducts and fomites), and aerosol (Meyer and Knudsen, 2001). The resulting loss of export markets from an FMD outbreak would bring severe economic consequences for producers of livestock, livestock products and livestock genetic material.

### 3.1. Outbreak scenario

EuFMDiS-FMD was configured for a study area comprising Austria, Croatia, Hungary, and Slovenia. The total livestock population in the study area was approximately 16 million animals spread across 316,443 herds. FMD was introduced into a large-scale commercial breeding pig holding in eastern Slovenia (latitude 46.6618, longitude 15.8537), approximately 7 km from the Austrian border. Four of the 1104 pigs were infected after illegal feeding of swill that contained contaminated pork products sourced from a country where FMD is present. The outbreak began on the 1st of September and was detected and reported to the authorities 21 days later.

### 3.2. Study design

In this case study we compare two control policies for managing the outbreak, one based on stamping out of infected herds and one based on stamping out plus emergency ring vaccination. 1000 iterations of the outbreak scenario were run firstly with a control program consistent with EU guidelines (European Union, 2003) comprising movement restrictions, stamping out, surveillance and tracing, and secondly with the addition of suppressive ring vaccination. Selected control parameters are provided in Table 4. The simulation was configured to end at the completion of the control program, i.e., post-outbreak surveillance was not enabled. The maximum length of a simulation was set to 500 days.

**Table 4.** Selected control program parameter settings for the EuFMDiS-FMD case study

| Control Parameter | Value |
| --- | --- |
| national livestock standstill | 3 days |
| protection zone (PZ) | Circle of 3 km radius enclosing each IH |
| surveillance zone (SZ) | Circle of 10 km radius enclosing each IH |
| ratio of false positive reports of clinical disease to true positive | 2.34:1 (McLaws et al., 2007) |
| forward tracing window | 14 days |
| backward tracing window | 14 days |
| average time needed for a direct trace (days) | 1 to 2 days (species-dependent) |
| average time needed for an indirect trace (days) | 2 to 3 days (species-dependent) |
| effectiveness of direct tracing | 80% to 96% (species-dependent) |
| effectiveness of indirect tracing | 65% to 76% (species-dependent) |
| non-compliance with direct movement controls inside PZ | 0% to 2% (country-dependent) |
| non-compliance with direct movement controls inside SZ | 5% to 15% (country-dependent) |
| reduction of indirect movements inside PZ | 90% to 98% (country-dependent) |
| reduction of indirect movements inside SZ | 75% to 85% (country-dependent) |
| surveillance visit duration (days) | 0.28 to 0.73 (herd type-dependent) |
| number of surveillance resources (min, max) | 10 to 30 (per country) |
| number of culling resources (min, max) | 5 to 30 (per country) |
| number of disposal resources (min, max) | 5 to 30 (per country) |
| number of decontamination resources (min, max) | 10 to 50 (per country) |
| number of vaccination resources (min, max) | 10 to 50 (per country) |
| time to cull a herd (days) | 0.3 to 1.6 (herd type-dependent) |
| cost of culling an animal (€) | 16.8 to 38.4 (herd-type dependent) |
| time for disposal of a herd (days) | 0.5 to 1.5 (herd type-dependent) |
| cost of disposal of an animal | 19.3 to 117.4 (herd-type dependent) |
| time to decontaminate a holding (days) | 0.8 to 1.6 (herd type-dependent) |
| start of vaccination program | 7<sup>th</sup> day of the control program |
| time to vaccinate a herd (days) | 0.3 to 0.8 (herd type-dependent) |
| vaccination annulus radii (km) | 0, 3 |
| vaccination direction | outside-in |
| vaccine efficacy | 0.84 to 0.87 (species-dependent) |
| vaccinated herd types | all except backyard herds |

### 3.3. Statistical methods

The outcome variable distributions were not normally distributed, so were log transformed prior to t-test comparisons of the two control strategies. Means with 95% confidence intervals (CI) were calculated from the transformed distributions and then back (antilog) transformed (Table 5). Range statistics (5^th^ percentile, median, 95^th^ percentile and maximum), and shape statistics (skew and kurtosis) for selected (untransformed) outcome variables are presented in Table 6. RStudio (RStudio Team, 2020) was used to produce box and whisker plots for selected (untransformed) outcome variables. The box represents the 25–75 percentile range. The horizontal band within the box represents the median. The whiskers represent the 0–25 and 75–100 percentile ranges. The y-axes are presented logarithmically.

**Table 5.**
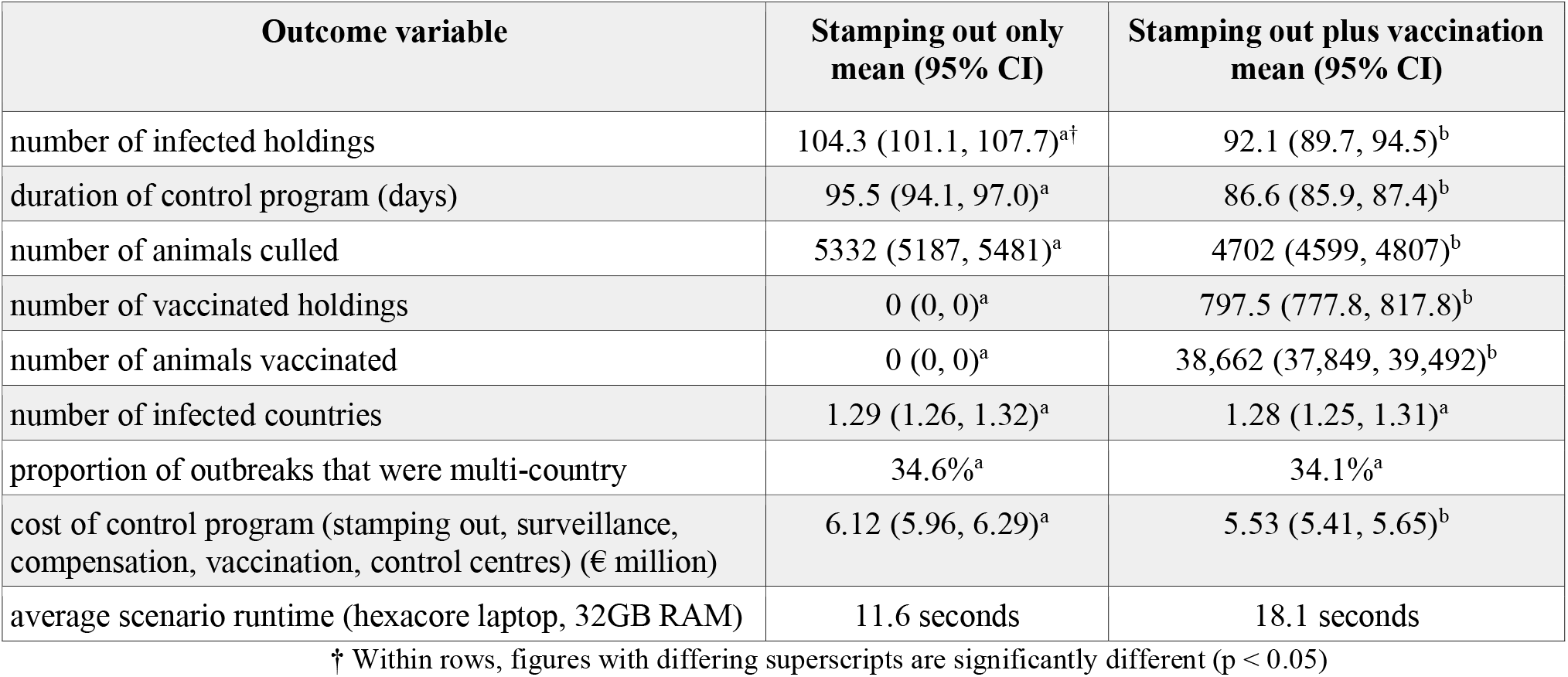
Selected outcomes of the EuFMDiS-FMD case study (across 1000 runs)

**Table 6.** Effect of vaccination on the distributions of selected outcome variables of the EuFMDiS-FMD case study

| Outcome variable | Distribution statistic | Stamping out only | Stamping out plus emergency vaccination |
| --- | --- | --- | --- |
| number of infected holdings | shape (skew, kurtosis) | (3.41, 20.32) | (1.94, 5.29) |
|  | range (5<sup>th</sup>, median, 95<sup>th</sup>, max) | (51, 96, 284, 990) | (50, 88, 196, 376) |
| duration of the control program | shape (skew, kurtosis) | (1.72, 3.94) | (1.12, 1.85) |
|  | range (5<sup>th</sup>, median, 95<sup>th</sup>, max) | (70, 91, 153, 262) | (71, 85, 112, 148) |
| number of animals culled | shape (skew, kurtosis) | (2.73, 12.68) | (1.89, 4.79) |
|  | range (5<sup>th</sup>, median, 95<sup>th</sup>, max) | (2951, 5051, 12844, 37,777) | (2871, 4455, 9184, 16,539) |

### 3.4. Results

Selected outcome variables are reported in Table 5 as means with 95% confidence interval. Figures 5 and 6 compare stamping out only and stamping out plus suppressive ring vaccination, with respect to the number of IHs and the duration of the control program. A screenshot of the 1000^th^ iteration of the stamping out plus vaccination scenario illustrates how EuFMDiS outcome variables are distributions (Figure 7). In this case, the median outbreak duration was 106 days (range 83 to 169 days) (blue graph), and the median number of IHs was 88 (range 32 to 376) (red graph).

**Figure 5.**
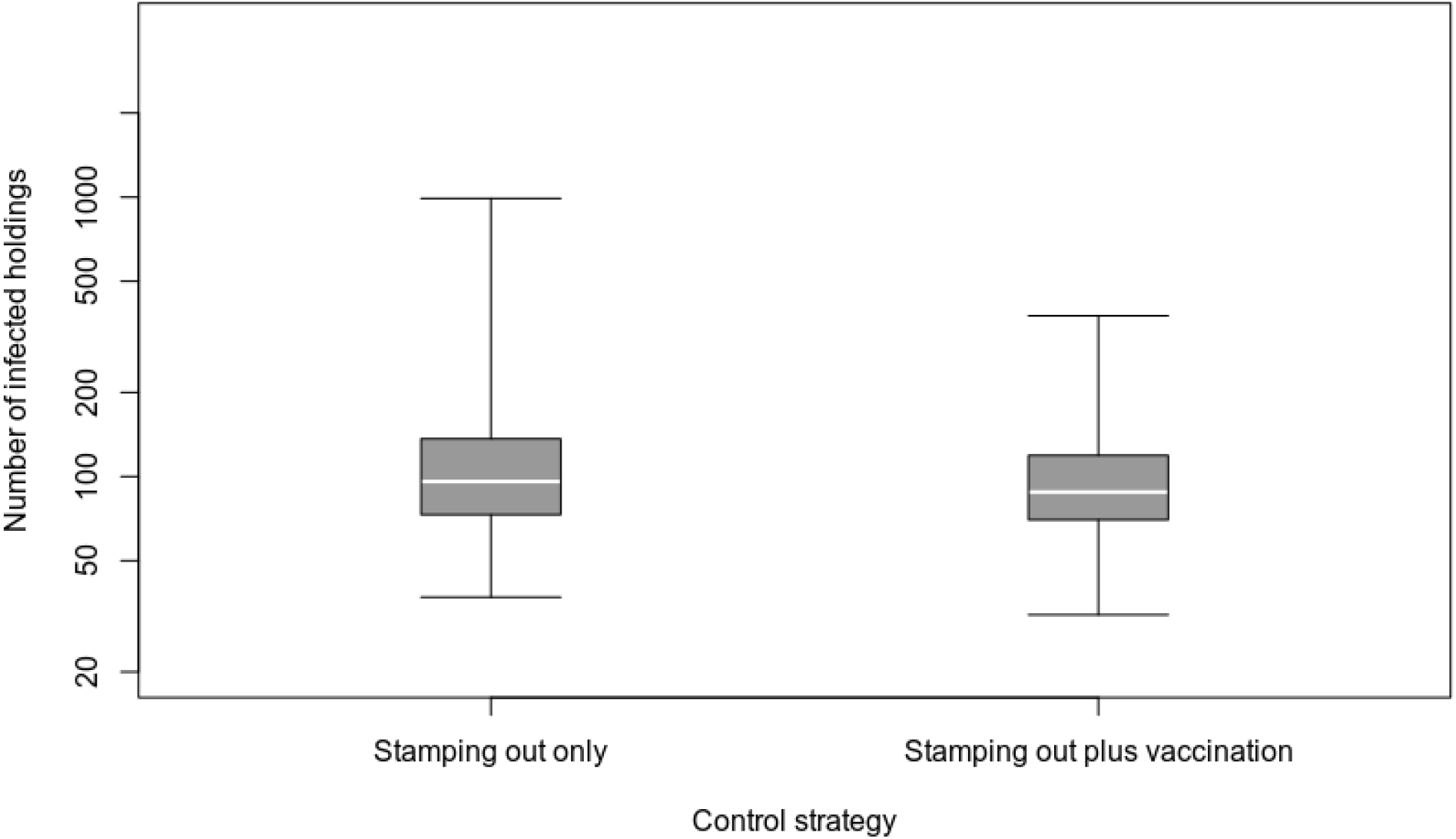
Effect of suppressive vaccination on the number of infected holdings.

**Figure 6.**
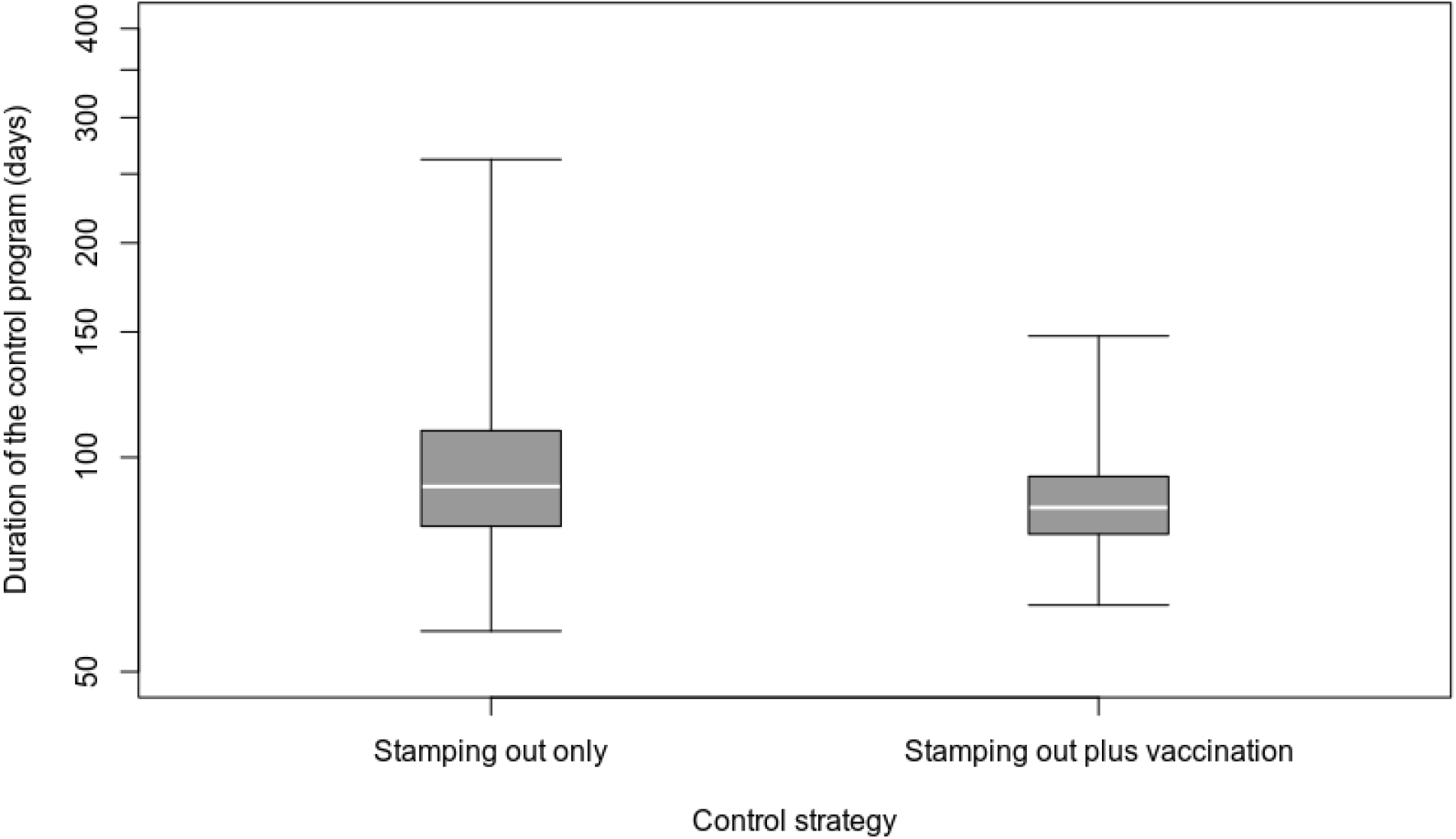
Effect of suppressive vaccination on the duration of the control program.

**Figure 7.**
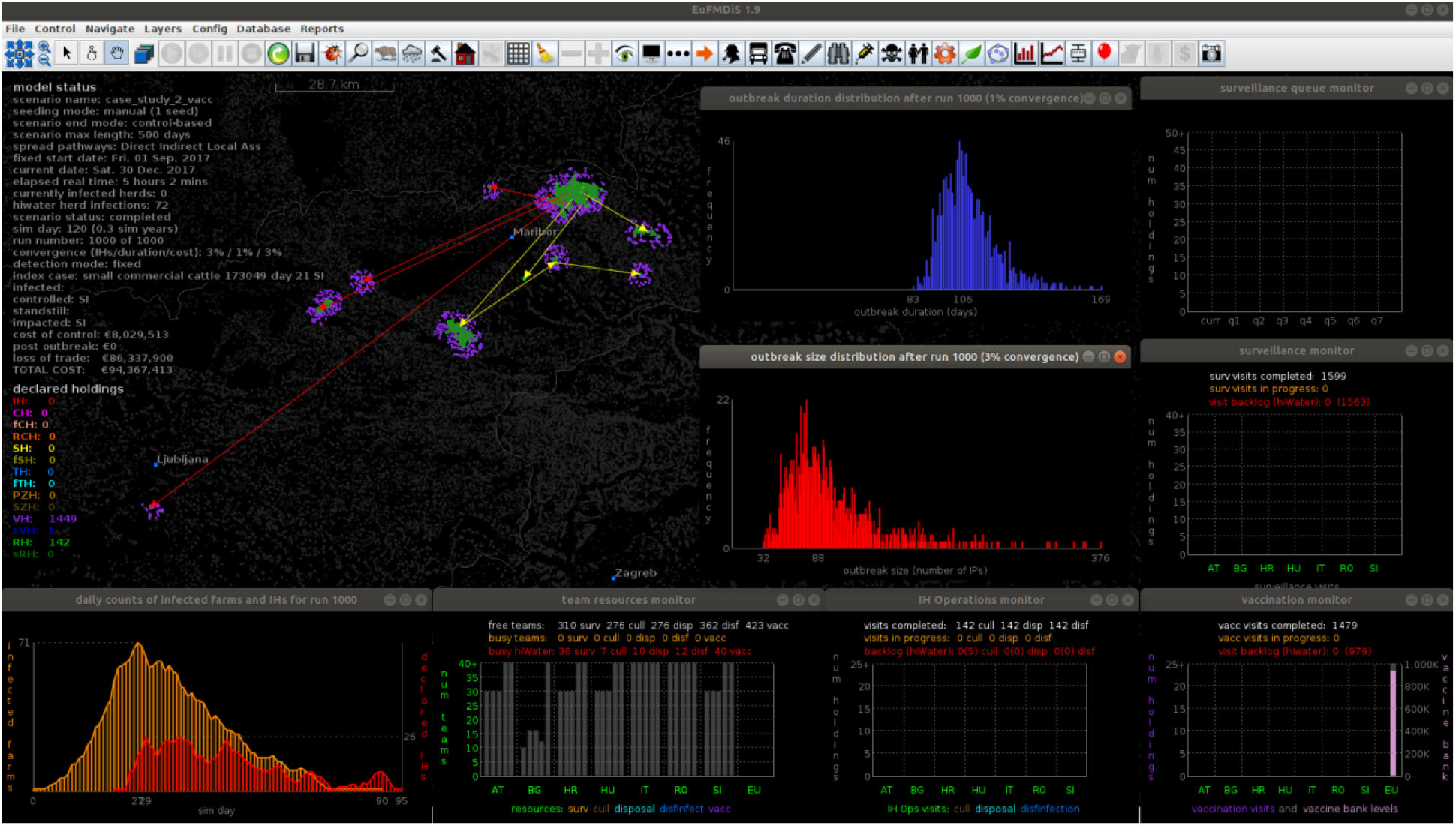
Screenshot of EuFMDiS-FMD after 1000 runs of the vaccination scenario in the case study.

### 3.5. Discussion

Augmenting the stamping out-only control policy with suppressive ring vaccination resulted in small, but statistically significant, improvements in the effectiveness and cost of control. On average, the duration of the control program was decreased by 8.9 days (approximately a 9% reduction), the number of IHs by 12.3 (12%), the number of culled animals by 630 (12%), and the cost of the control program by €590,350 (10%) (Table 5). Vaccination was effective in reducing outbreak variability and the likelihood of a large outbreak (Figures 5 and 6). Table 6 shows that vaccination reduced the ‘long tail’ of the outbreak with significant reductions in skew, kurtosis, and the upper range of the outcome distributions.

The case study involved a specific primary case and so it is unwise to generalise findings on control. EuFMDiS offers more general means for the user to seed outbreaks in time and space. For example, instead of always seeding the initial infection in the same large-scale commercial pig breeding holding in eastern Slovenia, each iteration of the outbreak could be seeded in a randomly chosen large-scale commercial breeding pig holding in Slovenia that has 350 or more pigs.

The case study did not include post-outbreak surveillance to support of proof-of-freedom and management strategies for vaccinated animals. Whilst these features are available in EuFMDiS, they were omitted from the case study in the interests of brevity. It is unwise to draw conclusions on the cost-effectiveness of vaccination in a control program without also considering whether vaccinated animals are destroyed or retained in the population, and the related implications on post-outbreak surveillance and mandatory OIE waiting periods prior to return to trade (Bradhurst et al., 2019).

## 4. DISCUSSION

In a cooperative multinational setting such as the EU with high levels of trade and travel between member states, there is increased risk of a transboundary animal disease silently crossing borders via movements of presymptomatic infectious livestock, contaminated livestock products or fomites. Livestock diseases can also be conveyed between countries via natural intermediary pathways such as wild animals, insect vectors or windborne aerosols. The nature and extent of transboundary animal disease outbreaks depends on many variables including livestock density, production systems, marketing systems, climate, pathogen specifics, surveillance regimes, and animal health resources. It can be challenging for national disease managers to plan and prepare for outbreaks in the face of spatiotemporal heterogeneity. Further challenges arise when policy formulation, response planning and resourcing, biosecurity standards, and animal welfare issues need to be considered in a close-knit multinational context.

Epidemiological models are increasingly employed as decision support tools to assist disease preparedness and planning. Such models are, however, typically funded by individual countries and geared towards national concerns and priorities. A transboundary model of livestock disease would be a useful tool for decision makers; however, national models are not always easily extended beyond borders. Details on livestock population, production systems and marketing systems of neighbouring countries are not always available, and it can be a challenge to capture heterogeneities in production systems, control policies, and response resourcing across multiple countries, in a single model.

The EuFMDiS transboundary disease modelling framework was developed in a collegiate manner by epidemiologists, disease modellers, and the animal health agencies of seven EU member states. It simulates the spread of TADs within and between countries and allows control policies to be enacted and resourced on per-country basis. It provides a sophisticated decision support tool that can be used to look at the risk of disease introduction, establishment and spread; evaluate control approaches in terms of effectiveness and costs; study resource management; and address post-outbreak management issues. EuFMDiS can operate in either single-country mode (where only within-country spread and control of disease is simulated), or multi-country mode (where both within-country and between-country spread, and control of disease is simulated).

EuFMDiS is a hybrid model in that it combines analytical and mechanistic modelling approaches. The spread of disease within a herd is represented mathematically while the spread of disease between herds is represented with a data-driven agent-based approach. Data-driven agent-based models are well suited to capturing heterogeneity, stochasticity, spatial relationships, seasonality, social systems, and policy nuances. A downside of data-driven modelling approaches is that they depend heavily on the quality of the underlying data and require expert parameterisation. Mathematical models, on the other hand, are simpler to parameterise but often assume that a population is homogeneous and well-mixed, and thus fail to capture heterogeneities. A novel feature of EuFMDiS is that it offers either data-driven or analytical (jump-diffusion) spread pathways between herds. The analytical spread pathways are useful when there is insufficient or unreliable data available to parameterise the data-driven pathways.

Complex epidemiological models such as EuFMDiS require skilled and trained personnel to use and interpret findings properly. EuFMD is addressing this through a comprehensive user manual and initiatives such as promoting an active user community and online training courses.

The test case disease for EuFMDiS development was FMD. Development is underway for other important TADs including ASF and CSF, and the incorporation of wild boar as a vector pathway (Bradhurst et al., 2020).

## ACKNOWLEDGEMENTS

The authors would like to thank the European Commission for the Control of FMD (EuFMD) for their unwavering support and funding of the EuFMDiS development project. The EuFMDiS modelling framework is a multi-country adaptation of the Australian AADIS modelling framework and the authors would like to acknowledge the Australian Department of Agriculture, Water and the Environment who funded the development of AADIS, and made it available to the EuFMD. The authors would also like to acknowledge and thank the state veterinary authorities of Austria, Bulgaria, Croatia, Hungary, Italy, Romania, and Slovenia. The development of the pilot EuFMDiS-FMD model would not have been possible without their support, and provision of data and parameter values.

## CONFLICT OF INTEREST STATEMENT

To the author’s best knowledge, there are no conflicts of interest to report.

## DATA AVAILABILITY STATEMENT

The data supporting the case study are available from the corresponding author upon reasonable request.

## ETHICS STATEMENT

The authors confirm that the ethical policies of the journal, as noted in the journal’s author guidelines, have been adhered to. No ethical approval was required as this article reports on the development of a new decision support modelling framework.

## Notes

### Competing Interest Statement

The authors have declared no competing interest.

